# Negative immune regulation contributes to disease tolerance in Drosophila

**DOI:** 10.1101/2021.09.23.461574

**Authors:** Arun Prakash, Katy M. Monteith, Pedro F. Vale

## Abstract

Disease tolerance is an infection phenotype where hosts show relatively high health despite harbouring elevated pathogen loads. Variation in the ability to reduce immunopathology may explain why some hosts can tolerate higher pathogen burdens with reduced pathology. Negative immune regulation would therefore appear to be a clear candidate for a mechanism underlying disease tolerance. Here, we examined how the negative regulation of the immune deficiency (IMD) pathway affects disease tolerance in *Drosophila melanogaster* when infected with four doses of the gram-negative bacterial pathogen *Pseudomonas entomophila*. We find that while flies unable to regulate the IMD response exhibited higher expression of antimicrobial peptides and lower bacterial loads as expected, this was not accompanied by a proportional reduction in mortality. Instead, UAS-RNAi knockdown of negative regulators of IMD (*pirk* and *caudal*) substantially increased the per-pathogen-mortality in both males and females across all tested infectious doses. Our results therefore highlight that in addition to regulating an efficient pathogen clearance response, negative regulators of IMD also contribute to disease tolerance.

## Introduction

To survive infection hosts must achieve a fine balance between clearing pathogens and avoiding immune-induced pathology by tightly regulating the induction and resolution of immune responses [1–3]. Understanding these distinct pathogen- or host-derived causes of tissue damage is important not only from a therapeutic perspective [4–6], but also offers important mechanistic explanations for the variable infection phenotypes that are often observed between individuals [7–9]. Individuals with low pathogen loads and relatively high health during infection are commonly described as ‘resistant’ because they are able to clear - or otherwise quantitatively reduce - the establishment and growth of pathogens within the host [7,8,10,11]. Other infection phenotypes are more difficult to explain as the consequence of resistance, such as the observation that some individuals exhibit low levels of pathology (measured as either mortality or physiological performance), despite harbouring relatively high pathogen loads [8,9,12,13].

Instead, tolerating higher pathogen loads is more likely to result from mechanisms that prevent, limit or repair tissue damage [14–17]. Previous work in *Drosophila* uncovered several candidate genes associated with disease tolerance, including many that do not have an obvious immune related function such as *CrebA* [18], *grainyhead and debris buster* [19], *dFOXO* [20,21]. Curiously, genes related to canonical immune pathways are conspicuously absent from these genome-wide screens of tolerance, even though inflammation is often an underlying cause of damage during infection, and we might therefore expect the regulation of inflammation to be a key mechanism underlying disease tolerance [2,8,13,22].

Other work, however, suggests that minimization of inflammation may be key to understanding tolerance. For example, pro-inflammatory cytokines tend to be associated with decreased tolerance in populations of house finches (*Haemorhous mexicanus*) infected with a bacterial pathogen *Mycoplasma gallisepticum*, which exhibit reduced cytokine expression compared with a less tolerant population [23]. Lower levels of circulating pro-inflammatory cytokines are also associated with increased tolerance of malaria [24]. In mice, treating *Mycobacterium tuberculosis* infection with the anti-inflammatory drug Ibuprofen resulted in marked improvements in infection tolerance [25], while negative regulation of inflammation-like responses permit greater tolerance of Drosophila C virus infection in fruit flies [22,26,27]. Negative regulation of immune responses would therefore appear to be a promising candidate for a general a mechanism that promotes disease tolerance [16,24,28,29].

In the present work, we investigate if the negative regulation of immune responses has any measurable effect on the phenotype of disease tolerance during systemic bacterial infection in Drosophila. We focused on the regulation of the immune deficiency (IMD) pathway (Fig. S1), one the best described immune signalling pathways in *Drosophila* [30–32]. Following infection with gram-negative bacteria, DAP-type (Diaminopimelic acid) peptidoglycans are recognised by pathogen-sensing receptors *PGRPs (LC* and *LE)* [33,34]. These receptors then recruit the adaptor molecule IMD and activate an intracellular signalling cascade that leads to the activation and translocation to the nucleus of a *NF-*_ĸ_*B relish*, inducing the transcription of a set of IMD-responsive antimicrobial peptides (AMPs)[32]. *Relish* also induces the transcription of negative regulators of IMD (*pirk, caudal, PGRPs-LB, LC, LE*,) which ensures an appropriate level of immune response, while avoiding immunopathology [35,36]. *Pirk*, for example, interferes with the interaction of *PGRP-LC* and *-LE* with the molecule IMD, and limits the activation of the IMD pathway [37–39]. Similarly, the homeobox protein *caudal* downregulates the expression of AMPs in flies [40] (Fig. S1). Taken together, the negative regulation of IMD-pathway ensures an appropriate level of immune response and enables individuals to reduce immune activity in response to reductions in bacterial numbers [35,38,41].

We tested whether disrupting optimal regulation of the IMD pathway can impact the ability of Drosophila to tolerate bacterial infection. We measured disease tolerance during systemic infection with the gram-negative bacterium *Pseudomonas entomophila* using *Drosophila* lines where the expression of specific components of the IMD-pathway were knocked down using UAS-RNAi, including the transcription factor-*relish*, its negative regulators *pirk* and *caudal*, and a major IMD-responsive AMP *Diptericin-B*.

## Materials and methods

### Experimental methods

To test the role of IMD negative regulation on systemic bacterial infection, we used fly lines where the UAS/Gal4 system was used for ubiquitous downregulation of target genes with RNA interference (RNAi): UAS-RNAi-*relish*, UAS-RNAi-*caudal* UAS-RNAi-*pirk* and UAS-RNAi-*diptericinB* (see Supplementary Methods and Table S1 for details about fly genotypes). We infected these flies with four inoculation doses of the gram-negative bacterium *Pseudomonas entomophila* [34] OD_600_ = 0.1, 0.05, 0.01 and 0.005, delivering approximately I) ∼650 cells (II) ∼1400 cells (III) ∼2000 cells (IV) ∼2900 cells, respectively). Following infection, we quantified the expression of three IMD-responsive AMPs namely, diptericin (Dpt), attacin-C AttC), cecropin-A1 (CecA1), by quantitative real-time polymerase chain reaction (qRT-PCR) and quantified the effects of downregulating each gene of interest on the mortality of each fly and the internal bacterial loads achieved during the infection. A detailed description of fly rearing conditions and bacterial culture protocols, infection protocols, and the measurement of survival and bacterial loads is given in Supplementary Methods

## Statistical analysis

All data and code are available at. https://doi.org/10.5281/zenodo.5564519.

### Gene expression

We analysed the gene expression data by first calculating the ΔCT value [42] for the expression of a gene of interest relative to the reference house-keeping gene rp49 [26]. We analysed, using ANOVA, the effects of the experimental treatment on the ΔCT values, indicating the difference in AMPs expression between knockdown and fully functional flies of each IMD-pathway fly line. We used the R package ‘ggplot2’ for graphics and data visualization (Wickham 2016; Bunn and Korpela, 2019).

### Bacterial load

We quantified differences in bacterial load following bacterial infection by analysing the bacterial measurement (log_10_) 8 hours following *P. entomophila* infection. We found that residuals of bacterial load data were normally distributed when tested with Shapiro-Wilks test and hence we used ANOVA and analysed the bacterial load data by fitting ‘knockdown’ (functional flies and IMD-pathway knockdown flies), ‘sex’ and ‘infection dose’ (4 different infectious doses of *P. entomophila*) as categorical fixed-effects, while ‘vials’ (replicates) as a random-effects for each of the fly lines (functional flies and IMD-pathway knockdown flies).

### Survival post-infection

We analysed the survival data after *P. entomophila* bacterial infection for all the IMD-pathway mutants using a mixed-effects Cox model using the R package ‘coxme’ [45] for both males and females respectively. In each case, we specified the model as: survival ∼ knockdown _*(functional and knockdown)*_ x sex _*(male and female)*_ x dose _*(4 concentrations of P. entomophila)*_ + (1|vial _*(n replicates)*_), with ‘knockdown’ and ‘sex’ as fixed effects, and vials as a random effect for all the fly lines (IMD-pathway mutants). We fit separate models for each knockdown fly line and functional (wild type) fly line, since they were assayed independently on different days. We estimated the impact of harbouring IMD-pathway knockdowns as the hazard ratio of IMD-knockdowns vs. fully functional fly lines. A hazard ratio significantly greater than one indicates higher risk of mortality in the IMD-mutants relative to fully functional individuals; hence, a significant impact on fly’s survival.

### Measuring disease tolerance

We calculated tolerance as fly survival relative to its bacterial load (lifespan/mean bacterial load, also known as the per pathogen mortality (hours survival^-1^/ CFU/fly), akin to the often used per-pathogen-pathogenicity. This is metric of tolerance is intuitive to interpret: relative to a given microbe load, does one line die more (less tolerant) or less (more tolerant) [46– 49]. We analysed the per-pathogen-mortality (PPM) as PPM ∼ knockdown _*(functional and knockdown)*_ x sex _*(male and female)*_ x dose _*(4 concentrations of P. entomophila)*_.

## Results

*Relish* knockdown flies had lower expression of IMD-regulated AMP expression (*AttC, CecA1* and *Dpt*) and higher internal loads of *P. entomophila*, as expected. Loss of negative regulators *caudal* and *pirk* resulted in higher AMP expression and lower bacterial loads (***Fig***. *1*, ***Table****-S3-4)*. These results were consistent at three other infection doses (***Fig***.*S2* for bacterial load across different infection doses).

Given our observation that knocking down negative regulators of IMD resulted in an increase in AMP expression and lower microbe loads, we would expect mortality in these flies to be lower than controls if variation in mortality was mainly explained by pathogen clearance. However, while mortality was higher in *relish* and *DiptericinB* knockdowns as expected under reduced resistance, we also observed higher mortality when knocking down the expression of both negative regulators, *pirk* and *caudal* (*Fig. S3-S5, Table S5 for survival curves and hazard ratios across different infection doses*). This occurred despite the increased expression of AMPs and their relatively lower bacterial load, which is suggestive of reduced tolerance.

To quantify the tolerance response of each fly line more precisely, we measured the per-pathogen-mortality (measured as fly survival relative to its bacterial load) of male and female RNAi-lines of the IMD-pathway (*relish, caudal, pirk* and *DptB*), across four infectious doses (ranging from ∼650 bacterial cells infection dose to leading up to ∼2900 cells). Overall, we found that flies with UAS-RNAi-mediated knockdown in expression any of the genes of interest showed an increase in the per-pathogen-mortality, for all tested infection doses, and without substantial differences between sexes (Fig. 2, Table 1). In flies with reduced *caudal* expression, we detected significant sex-by-knockdown interaction, driven by smaller effects of UAS-RNAi of *caudal* in males at higher doses (Table 1).

**Table 1.**
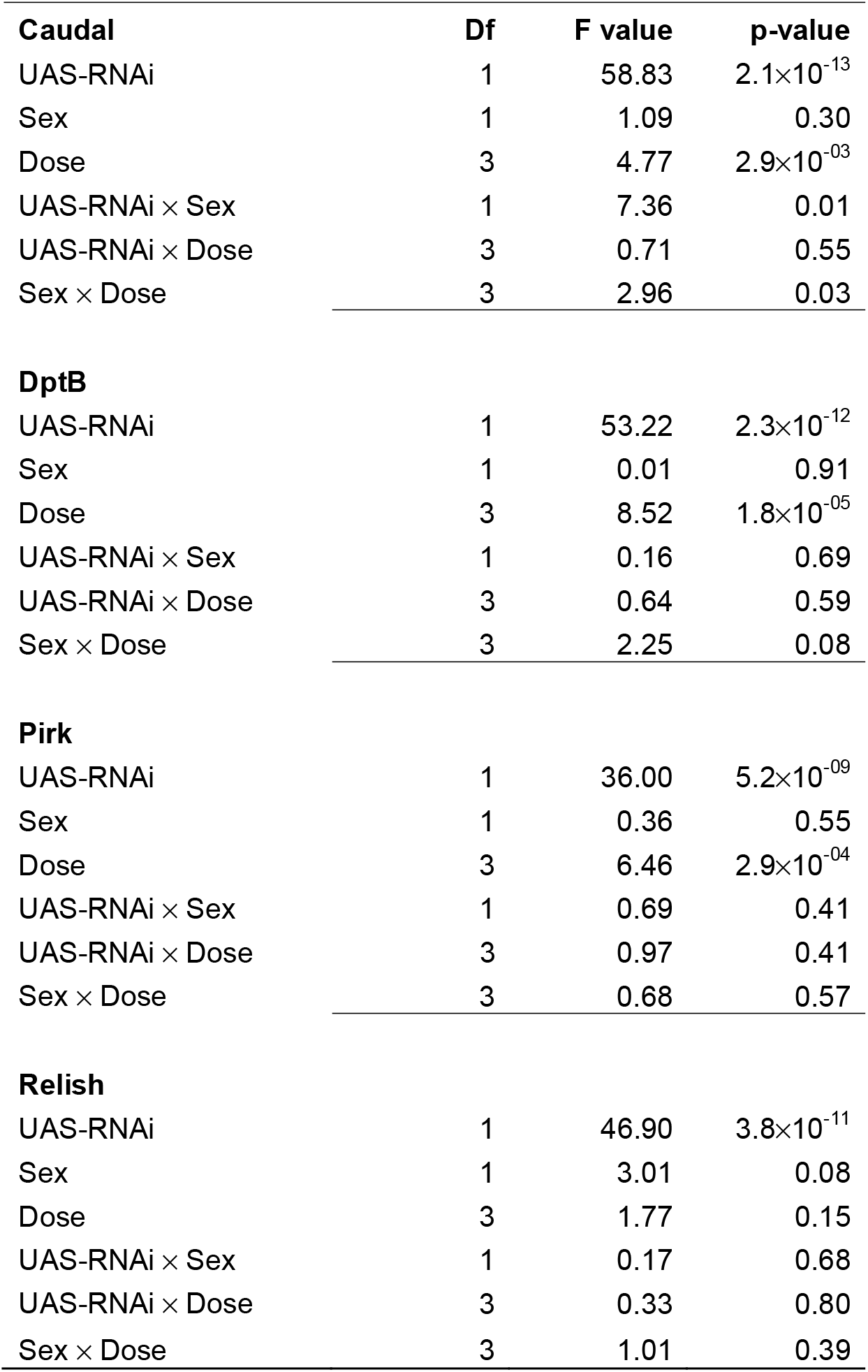
ANOVA testing the effects on the per-pathogen-mortality of RNAi knockdown of the expression of each gene of interest, fly sex and infection dose of *P. entomophila*. Details of statistical analysis are given in the methods section.

| **Caudal** | **Df** | **F value** | **p-value** |
| --- | --- | --- | --- |
| UAS-RNAi | 1 | 58.83 | $2.1 \times 10^{-13}$ |
| Sex | 1 | 1.09 | 0.30 |
| Dose | 3 | 4.77 | $2.9 \times 10^{-03}$ |
| UAS-RNAi × Sex | 1 | 7.36 | 0.01 |
| UAS-RNAi × Dose | 3 | 0.71 | 0.55 |
| Sex × Dose | 3 | 2.96 | 0.03 |

| **DptB** |  |  |  |
| --- | --- | --- | --- |
| UAS-RNAi | 1 | 53.22 | $2.3 \times 10^{-12}$ |
| Sex | 1 | 0.01 | 0.91 |
| Dose | 3 | 8.52 | $1.8 \times 10^{-05}$ |
| UAS-RNAi × Sex | 1 | 0.16 | 0.69 |
| UAS-RNAi × Dose | 3 | 0.64 | 0.59 |
| Sex × Dose | 3 | 2.25 | 0.08 |

| **Pirk** |  |  |  |
| --- | --- | --- | --- |
| UAS-RNAi | 1 | 36.00 | $5.2 \times 10^{-09}$ |
| Sex | 1 | 0.36 | 0.55 |
| Dose | 3 | 6.46 | $2.9 \times 10^{-04}$ |
| UAS-RNAi × Sex | 1 | 0.69 | 0.41 |
| UAS-RNAi × Dose | 3 | 0.97 | 0.41 |
| Sex × Dose | 3 | 0.68 | 0.57 |

| **Relish** |  |  |  |
| --- | --- | --- | --- |
| UAS-RNAi | 1 | 46.90 | $3.8 \times 10^{-11}$ |
| Sex | 1 | 3.01 | 0.08 |
| Dose | 3 | 1.77 | 0.15 |
| UAS-RNAi × Sex | 1 | 0.17 | 0.68 |
| UAS-RNAi × Dose | 3 | 0.33 | 0.80 |
| Sex × Dose | 3 | 1.01 | 0.39 |

## Discussion

Compared to the mechanisms of immune clearance our knowledge of the mechanisms underlying increased tolerance remains incomplete. We predicted that knocking down the expression of negative immune regulators would reduce the ability to flies to tolerate infection with *P. entomophilia*, reflected as an increase in the per-pathogen-mortality. While this was observed, we also observed higher per-pathogen mortality in *relish* and *dptB* knockdown flies. However, the underlying cause of this increase is likely to differ between lines (Figure S6). Flies with reduced expression of *relish* or *DptB* showed higher per-pathogen mortality, but internal loads in these flies were at least an order of magnitude higher than controls because of disrupted AMP production (Fig 1). A higher per-pathogen-mortality in this case may therefore reflect feedbacks between increased pathogen growth and decreased ability to limit damage when microbe loads get too high (that is, the per-pathogen mortality becomes higher in flies unable to control pathogen loads in the first place).

**Figure 1:**
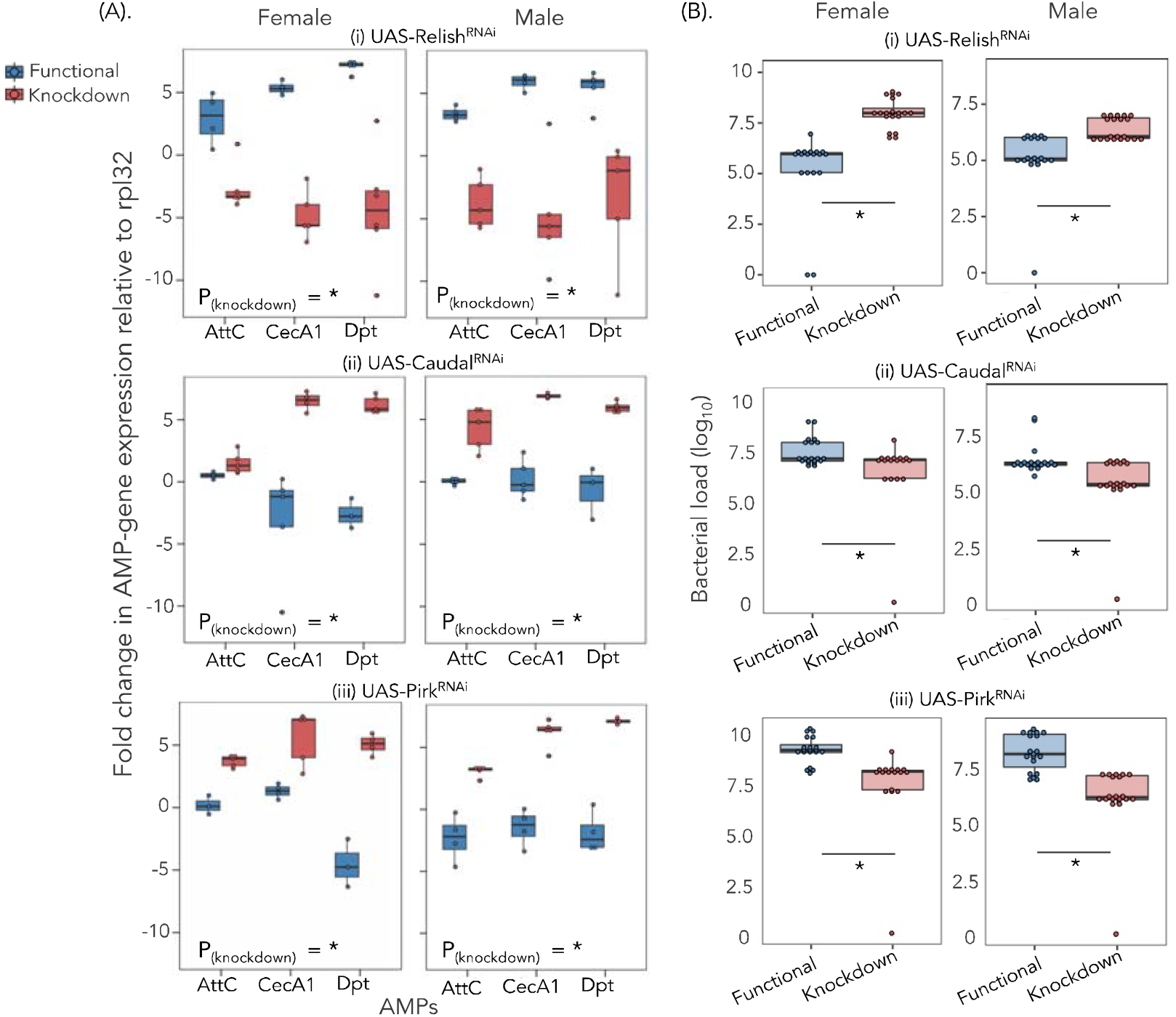
**(A)** Quantification of major IMD-responsive AMPs by qRT-PCR (n=15-21 flies/sex/treatment) for IMD-pathway mutants and functional flies (males and females) around 8 hours following *P. entomophila* systemic infection (0.05 OD or ∼2000 bacterial cells/μl). Each data point represents pooled data of 3 flies for each infection treatment, fly line and sex **(B)** internal bacterial load after 8 hours following infection (n=16-24 flies/sex/treatment). Asterisks inside the panels indicate significant differences in AMP expression (that is, impact of knocking down on overall AMPs measured) and bacterial load between knockdown flies and flies with fully intact IMD functioning (p<0.05). The error bars represent standard error.

**Figure 2:**
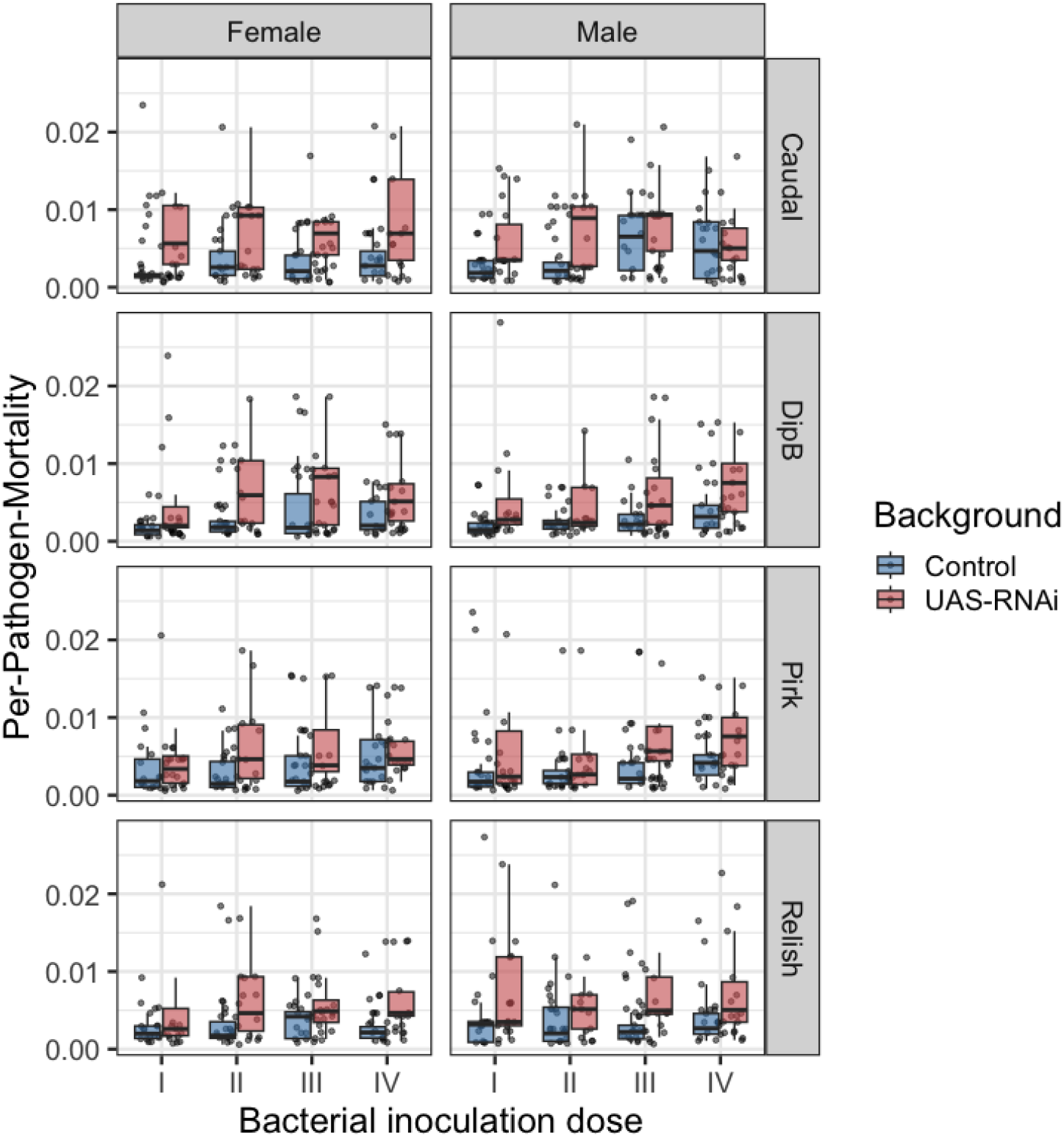
Disease tolerance in IMD-pathway mutants and fully functional flies. The relationship between host (fly) survival and bacterial load of 4 infection doses [that is, (I) ∼650 cells (II) ∼1400 cells (III) ∼2000 cells (IV) ∼2900 cells] of *P. entomophila*, analysing using point tolerance 8 hours following *P. entomophila* infection for **(A)** male and **(B)** female flies. Each data represents ‘point tolerance’, the ratio estimates calculated from median lifespan and average bacterial load for IMD-pathway knockdown and flies with fully functional IMD. Asterisks inside the panel indicates significant difference in disease tolerance between knockdown flies and with fully intact IMD functioning (p<0.05) across different infection doses. The error bars represent standard error.

By contrast, UAS-RNAi of the negative regulators *caudal* and *pirk* caused higher per-pathogen mortality despite these flies being able to achieve lower pathogen loads compared to functional control flies (Fig. 1). This is indicative of reduced tolerance, as flies experienced greater mortality despite harbouring fewer pathogens. Our results therefore highlight that the negative regulators *caudal* and *pirk* are important not only in regulating pathogen clearance by the IMD pathway, but by affecting the mortality of flies independently of their microbe loads, negative regulators of IMD would fit the functional definition of a tolerance mechanisms [9,13,21].

This increased mortality of flies lacking negative regulators of IMD such as *pirk* and *caudal* has been observed previously [38,40,50] and highlights that mechanisms other than pathogen clearance actively contribute to the variation in mortality during systemic bacterial infection. One likely cause of flies lacking negative IMD regulation still succumb to death despite reducing pathogen burden is immunopathology due to continued immune activation. In a similar example of negative immune regulation affecting disease tolerance, *G9a*, a negative regulator of Jak/Stat, was identified as being important in tolerating *Drosophila C Virus* infections by reducing immunopathology during the antiviral immunity [22]. During bacterial infections, *PGRP-LB* functions as a negative regulator of the IMD pathway by regulating the IMD-responsive AMP *Diptericin* [51], and it has been shown that *PGRP-LB* mutant flies experience sex-differences in survival despite undetectable differences in their bacterial loads [52], which is consistent with an effect on tolerance. We note that in the current experiment we did not detect substantial sex differences in the per-pathogen-mortality.

Recognising the role of negative immune regulators in disease tolerance phenotype opens up the possibility of using them as therapeutic targets to improve host health [4,5,14,53]. Future work may also investigate if natural genetic variation in disease tolerance can be explained by variation in negative immune regulators [27], offering a more complete mechanistic understanding of host heterogeneity in infection outcomes.

## Acknowledgments

We thank Angela Reid, Lucinda Rowe, James King and Alison Fulton for help with media preparation. We acknowledge funding and support from the Branco Weiss fellowship and a Chancellor’s Fellowship to PFV; a Darwin Trust PhD studentship to AP from the School of Biological Sciences, The University of Edinburgh. For the purpose of open access, the author has applied a Creative Commons Attribution (CC BY) licence to any Author Accepted Manuscript version arising from this submission.

