## Supplementary Methods for "Negative immune regulation contributes to disease tolerance in Drosophila"

### Electronic Supplementary Material

### Supplementary Methods

Fly strains and husbandry: We used the following GAL4/UAS fly lines for whole-fly silencing of target genes with RNA interference (RNAi) : UAS-RNAi-*relish* (Bloomington stock# 33661), UAS-RNAi-*caudal* (BS#57546), UAS-RNAi-*pirk* (BS#67011) and UAS-RNAi-*diptericinB* (BS#28975) (see Supplementary Information Table S1 for details about fly genotypes). All fly lines were obtained from the Bloomington Drosophila stock centre. Each of the transgenic fly line contains a homologous hairpin under the control of the Upstream Activating Sequence (UAS-RNAi) to a Gal4 temperature-sensitive transcriptional activator (Bloomington stock# 67065). This fly line drives the expression of Gal4 in the entire body of fly and contains a temperature sensitive Gal80, which inhibits Gal4 at the permissive temperature (25℃) where all the relevant genes are expressed normally. We used the driver fly line as a functional fly line (control). All fly stocks were reared in plastic vials (12 ml) on a standard Lewis medium (fly food) at 25±2 ̊C with a 12-hour light: dark on 14 days cycle (by transferring flies into new food vials). All the crosses were made at 25℃ and after eclosion, offspring were transferred to agar vials (12 flies per vial) at the non-permissive temperature (29℃) for two days to facilitate silencing of the candidate genes until experimental day.

***Table S1:*** Fly strains used in this study

| ***Fly line*** | ***Bloomington stock #*** | ***Genotype*** |
| --- | --- | --- |
| UAS-*relish^RNAi^* | *33661* | *y[1]sc[*]v[1]; P{y[+t7.7]v[+t1.8]=TRiP.HMS00070}attP2* |
| UAS-*caudal^RNAi^* | *57546* | *y[1]sc[*]v[1];P{y[+t7.7] v[+t1.8]=TRiP.HMC04863}attP40*), |
| UAS-*pirk^RNAi^* | *67011* | *y[1]sc[*]v[1];P{y[+t7.7] v[+t1.8]=TRiP.HMS05477}attP40* |
| UAS-*Diptericin*B*^RNAi^* | *28975* | *y[1]v[1];P{y[+t7.7] v[+t1.8]=TRiP.HM05186}attP2* |
| (*UAS^RNAi^*) ubiquitous driver | *67065* | w*;P{UAS-3xFLAG.dCas9.VPR}attP40, P{tubP-GAL80^ts^}10; P{tubP-GAL4}LL7/TM6B, Tb^1^ |

### Bacterial culture preparation: The gram-negative bacterium Pseudomonas entomophila has a broad host range, infecting insects, nematodes, plants, and higher vertebrates [1]. In flies, P. entomophila infection results in pathology to the gut epithelium during systemic infections, eventually causing death [1–3]. To obtain pure cultures, frozen bacterial stock cultures (-70℃) were streaked onto fresh LB agar plates and single colonies were inoculated into 15ml LB broth and incubated overnight at 37°C with shaking at 120rpm (revolutions per minute). The overnight cultures were diluted 1: 100 into fresh LB broth of 15ml and incubated again at 37°C with shaking at 120rpm. At the mid-log phase (optical density OD_600_ = 0.75), we harvested the bacterial cells by centrifugation at 5000 rpm for 5 min at 4°C and re-suspended the bacterial pellet in 1XPBS (phosphate buffer saline) solution. The final inoculum was adjusted to OD_600_ = 1.0, and this was the starting bacterial inoculum used to prepare 4 different dilutions of infection doses (OD_600_ = 0.1, 0.05, 0.01 and 0.005) delivering approximately I) ~650 cells (II) ~1400 cells (III) ~2000 cells (IV) ~2900 cells, respectively) during systemic infection.

### Gene expression: We quantified the expression of three IMD-responsive AMPs namely, diptericin (Dpt), attacin-C AttC), cecropin-A1 (CecA1), by quantitative real-time polymerase chain reaction (qRT-PCR). We randomly selected a subset of IMD-pathway mutants and functional flies both males and females at 8 hours following infection for RNA extraction, we included 15-21 flies that is, 3 flies pooled together for each treatment (Knockdown and functional flies) and sexes. We first homogenised whole flies (3 in group) using sterile micro-pestles into 1.5μl microcentrifuge tubes containing 80μL of TRIzol reagent (Invitrogen, Life Technologies). The tubes containing the fly homogenate were kept frozen at -70°C until RNA extraction. We performed mRNA extractions using the standard phenol-chloroform method and included a DNase treatment (Ambion, Life Technologies). We confirmed the purity of eluted samples using a Nanodrop 1000 Spectrophotometer (version 3.8.1) before proceeding with RT (reverse transcription). The cDNA was synthesized from 2μL of the eluted RNA using M-MLV reverse transcriptase (Promega) and random hexamer primers, and then diluted 1:1 in nuclease free water. We then performed quantitative RT-PCR (qRT-PCR) on an Applied Biosystems StepOnePlus machine using Fast SYBR Green Master Mix (Invitrogen Ltd.) using a 10μL reaction containing 1.5L of 1:1 diluted cDNA, 5μL of Fast SYBR Green Master Mix a 3.5μL of a primer stock containing both forward and reverse primer at 1μM suspended in nuclease free water (final reaction concentration of each primer being 0.35μM). For each cDNA sample, we performed two technical replicates for each set of primers and the average threshold cycle (Ct) was used for the analysis. We obtained the AMP primers from Sigma-Aldrich Ltd (see supplementary Table SI-2 for information about primers). We optimised the annealing temperature (Tₐ) and the efficiency (Eff) of the Dpt primer pair was calculated by 10-fold serial dilution of a target template (each dilution was assayed in duplicate); Dpt: Tₐ= 59 °C, Eff= 102%; AttC: Tₐ= 60 °C, Eff= 94%; CecA1: Tₐ= 59.5 °C, Eff= 109%.

***Table S2****:* Primers used in this study

|  | ***Obtained from*** | ***AMPs*** | ***Sequence*** |
| --- | --- | --- | --- |
| *1.* | [4] | *Dpt*_Forward: | 5’ GACGCCACGAGATTGGACTG 3’ |
|  |  | *Dpt*_Rverese | 5’ CCCACTTTCCAGCTCGGTTC 3’ |
|  |  | *AttC*_Forward: | 5’ TGCCCGATTGGACCTAAGC 3’ |
|  |  | *AttC*_Reverse: | 5’ GCGTATGGGTTTTGGTCAGTTC 3’ |
| *2.* | [5] | *CecA1*_Forward | 5’ TCTTCGTTTTCGTCGCTCTCA 3’, |
|  |  | *CecA1*_Reverse | 5’ ATTCCCAGTCCCTGGATT GTG 3’. |
| *3.* | [6] | *Rp49*_Forward: | 5’ ATGCTAAGCTGTCGCACAAATG 3’ |
|  |  | *Rp49*_Reverse: | 5’ GTTCGATCCGTAACCGATGT 3’ |

*Fly survival following bacterial infection:* To test the impact of *Pseudomonas entomophila* infection on fly survival, we challenged experimental flies to one of four bacterial cultures with increasing bacterial concentrations: (I) ~650 cells (II) ~1400 cells (III) ~2000 cells (IV) ~2900 cells/µl of *P. entomophila* culture, obtained by serial dilution of *P. entomophila* culture in exponential growth phase (see *bacterial culture preparation* above) . Flies (n=16-24 flies /sex/infection dose/fly line) were infected by intra-thoracic pricking [7] with a tungsten needle immersed in bacterial suspension under minimal CO_2_ anaesthesia. The effect of injury caused by pricking was controlled by including mock infections performed using a needle dipped in 1xPBS (phosphate buffer saline). We recorded the mortality for 10 days post-infection every 3 hours (±0.5 hours) for the first 2 days (between 6 am and 11 pm) and every 12 hours for the following 7 days.

*Quantification of bacterial load following bacterial infection:* During the survival assay, flies were randomly sampled (16-24 flies/sex/infection dose/fly line) and the bacterial load was quantified around 8 hours following infection. Initially, flies (group of 3) were transferred to a 1.5ml microcentrifuge tubes and surface sterilized by adding 70% ethanol for 30-60 seconds to ensure we counted only colony forming units (CFUs) inside the flies. Ethanol was then removed by briefly washing flies twice with sterile distilled water. To ensure that this method was efficient in surface cleaning flies, we plated 4µl of the second wash on LB agar (from random flies) ensuring that the flies are free from surface bacteria (that is, no viable CFUs). We then placed in 1.5ml Eppendorf tubes and added 100µl of LB broth to each tube and the flies were thoroughly homogenized using a motorized pestle. We immediately plated this homogenate in serial dilutions ranging from 100 to 10^-6^ using 1xPBS (phosphate buffer) on LB agar plates and incubated at 30°C for about 18 hours after which viable colonies were counted manually [8].

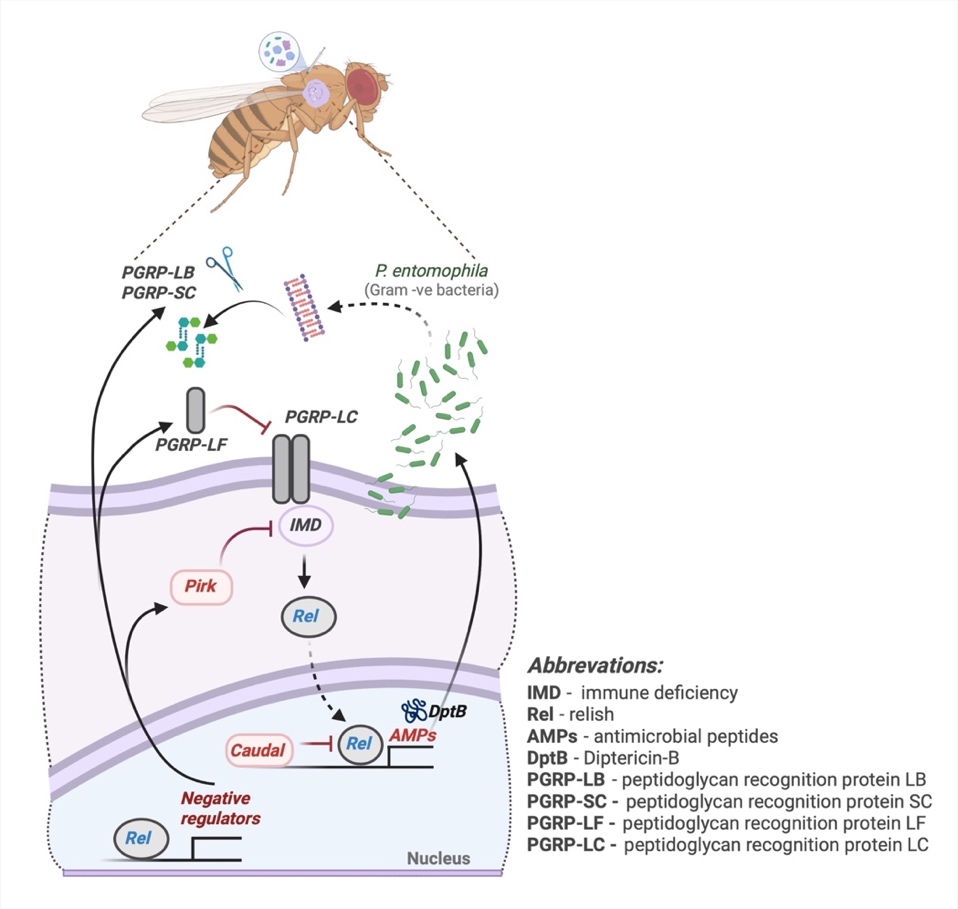

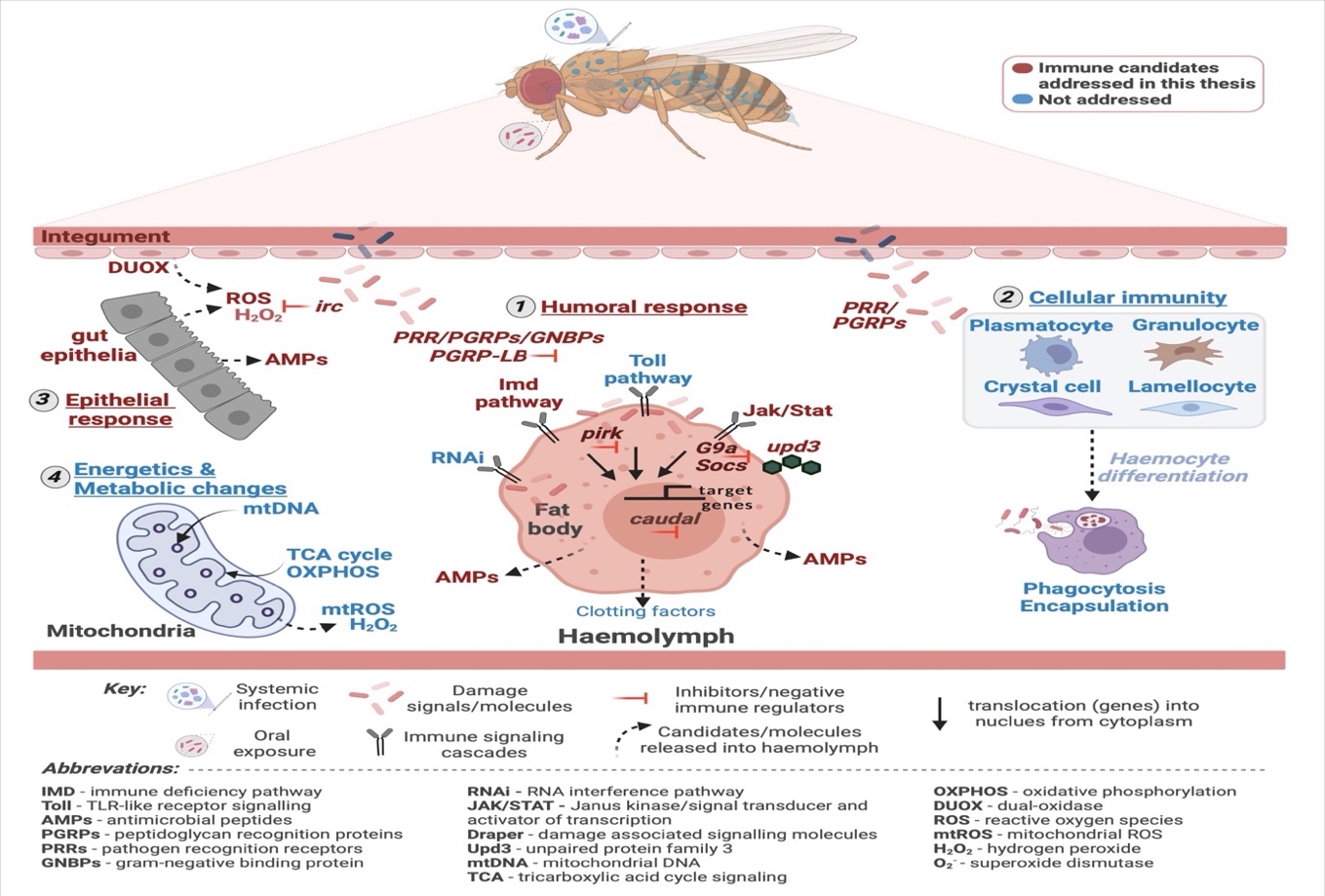

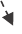

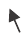
Figure S1: A simplified version of *Drosophila* IMD (immune deficiency) pathway activated by the DAP type pedptidoglycan following the gram-negative bacterial infection (*Pseudomonas entomophila*)*.* These receptors then recruit the adaptor molecule Imd and activate the intracellular signalling cascade that leads to the activation of a NF-ĸB *relish*. Upon activation, relish translocates to the nucleus and induces transcription of a set of cocktail of IMD-responsive AMPs such as *Diptericins, Attacins & Cecropins*, minutes after pathogen recognition. The negative regulators of IMD such as *pirk, caudal, PGRPs-LB* ensure an appropriate level of immune response, while avoiding immunopathology

***Figure S2*.** Bacterial loads around 24 hours after different concentrations of systemic *P. entomophila* infection for female flies for IMD-pathway mutants **(i)** *UAS-Relish^RNAi^* **(ii)** *UAS-Caudal^RNAi^* **(iii)** *UAS-Pirk^RNAi^* **(iv)** *UAS-DptB^RNAi^.*

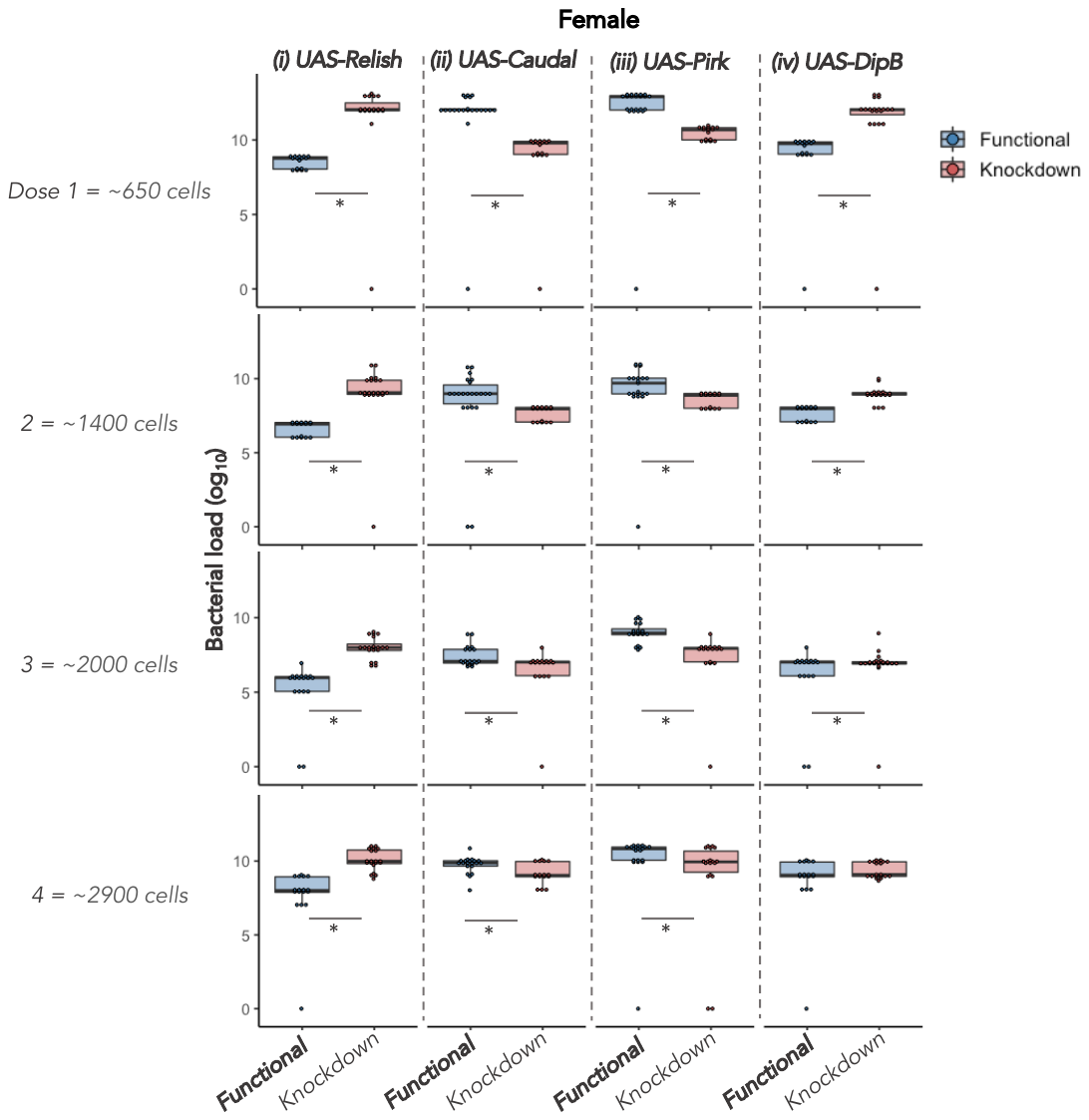

***Figure S3.*** Survival curves of female flies of different IMD-pathway mutants (n=16-24 flies/treatment/sex/fly line) **(a)** *UAS-Relish^RNAi^* **(b)** *UAS-Caudal^RNAi^* **(c)** *UAS-Pirk^RNAi^* **(d)** *UAS-DptB^RNAi^* during 4 concentrations of systemic *Pseudomonas entomophila* infection ranging between (1) ~650 cells (2) ~1400 cells (3) ~2000 cells (4) ~2900 cells/µl of *P. entomophila* culture. Since all 4 infection concentrations were performed simultaneously for each of the fly line, we used a single sham/mock-controls (SI) across all 4 different infection doses, (I) represents infection treatment.

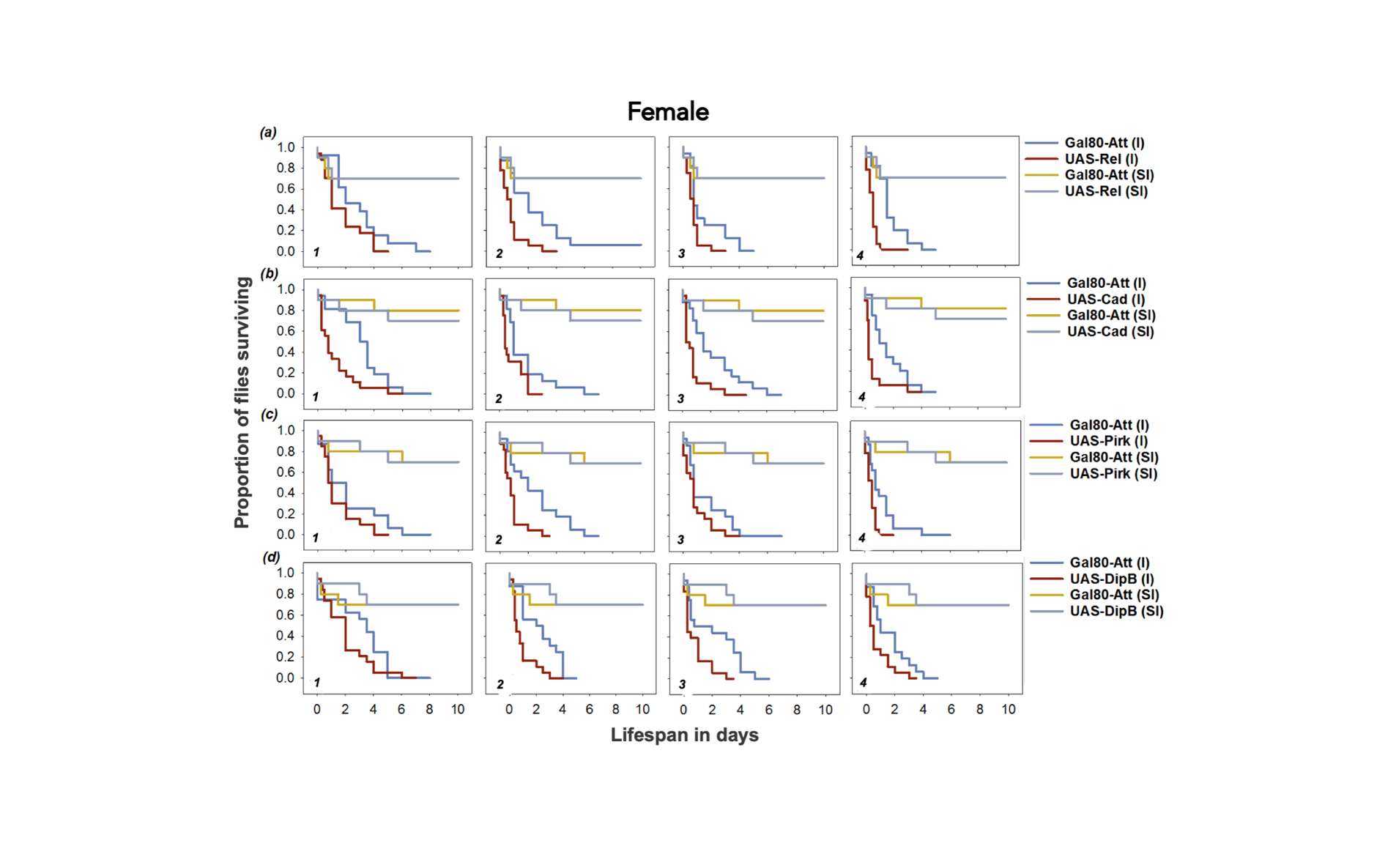

***Figure S4.*** Survival curves of male flies of different IMD-pathway mutants (n=16-24 flies/treatment/sex/fly line) **(a)** *UAS-Relish^RNAi^* **(b)** *UAS-Caudal^RNAi^* **(c)** *UAS-Pirk^RNAi^* **(d)** *UAS-DptB^RNAi^* during 4 concentrations of systemic *Pseudomonas entomophila* infection ranging between (1) ~650 cells (2) ~1400 cells (3) ~2000 cells (4) ~2900 cells/µl of *P. entomophila* culture. Since all 4 infection concentrations were performed simultaneously for each of the fly line, we used a single sham/mock-controls (SI) across all 4 different infection doses, (I) represents infection treatment.

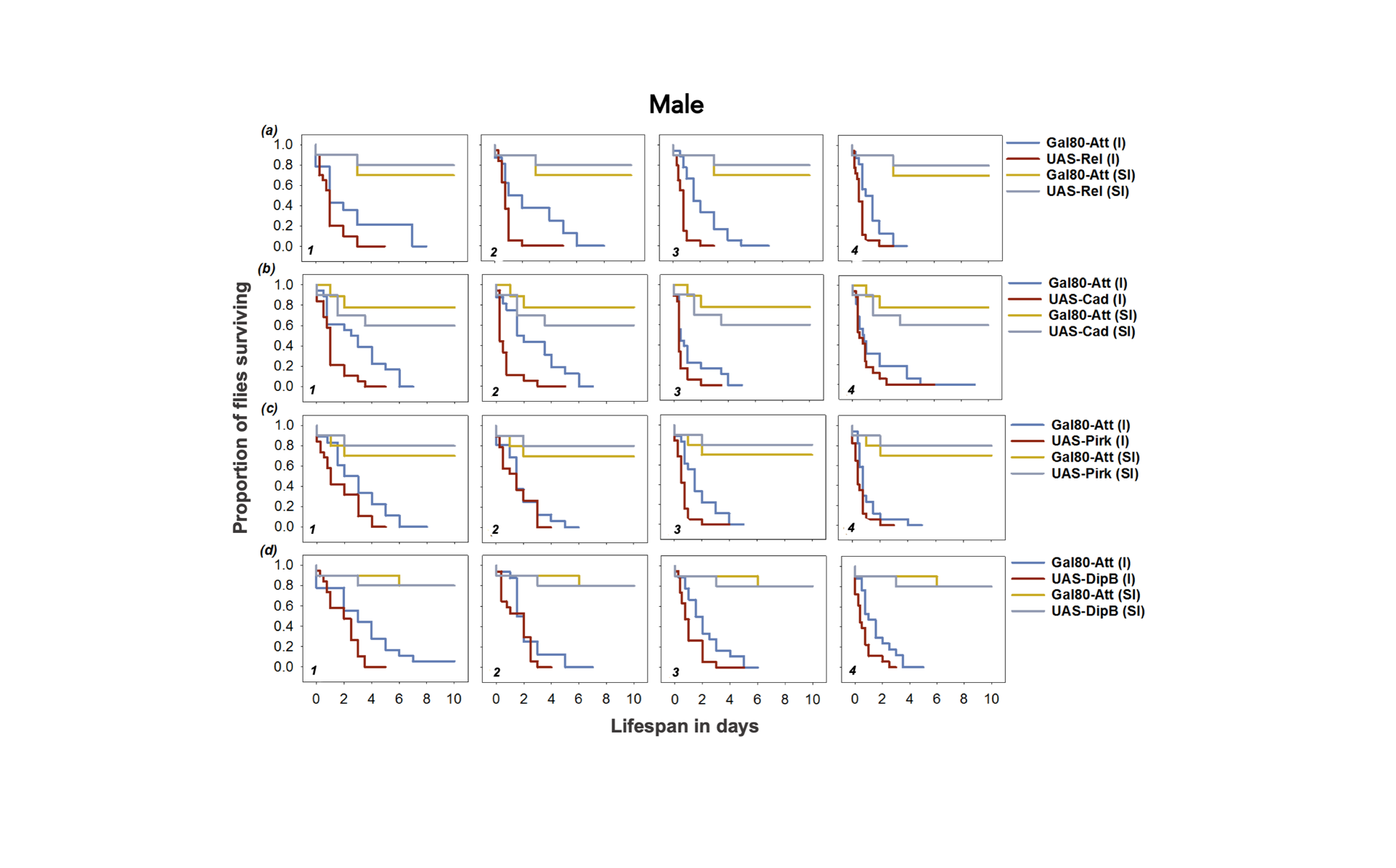

***Figure S5.*** The estimated hazard ratios calculated from survival curves (***Fig.*** *S1* *and* *S2*) for flies with fully functional immune system and IMD-pathway mutants for both Sexes. The different IMD*-*pathway mutants include **(a)** *UAS-Relish^RNAi^* **(b)** *UAS-Caudal^RNAi^* **(c)** *UAS-Pirk^RNAi^* **(d)** *UAS-DptB^RNAi^.* We used 4 different concentrations of systemic *P. entomophila* infection*.* A greater hazard ratio >1 shown as horizontal dashed grey lines indicates higher susceptibility of IMD*-*mutant flies than the functional flies to *P. entomophila* infection.

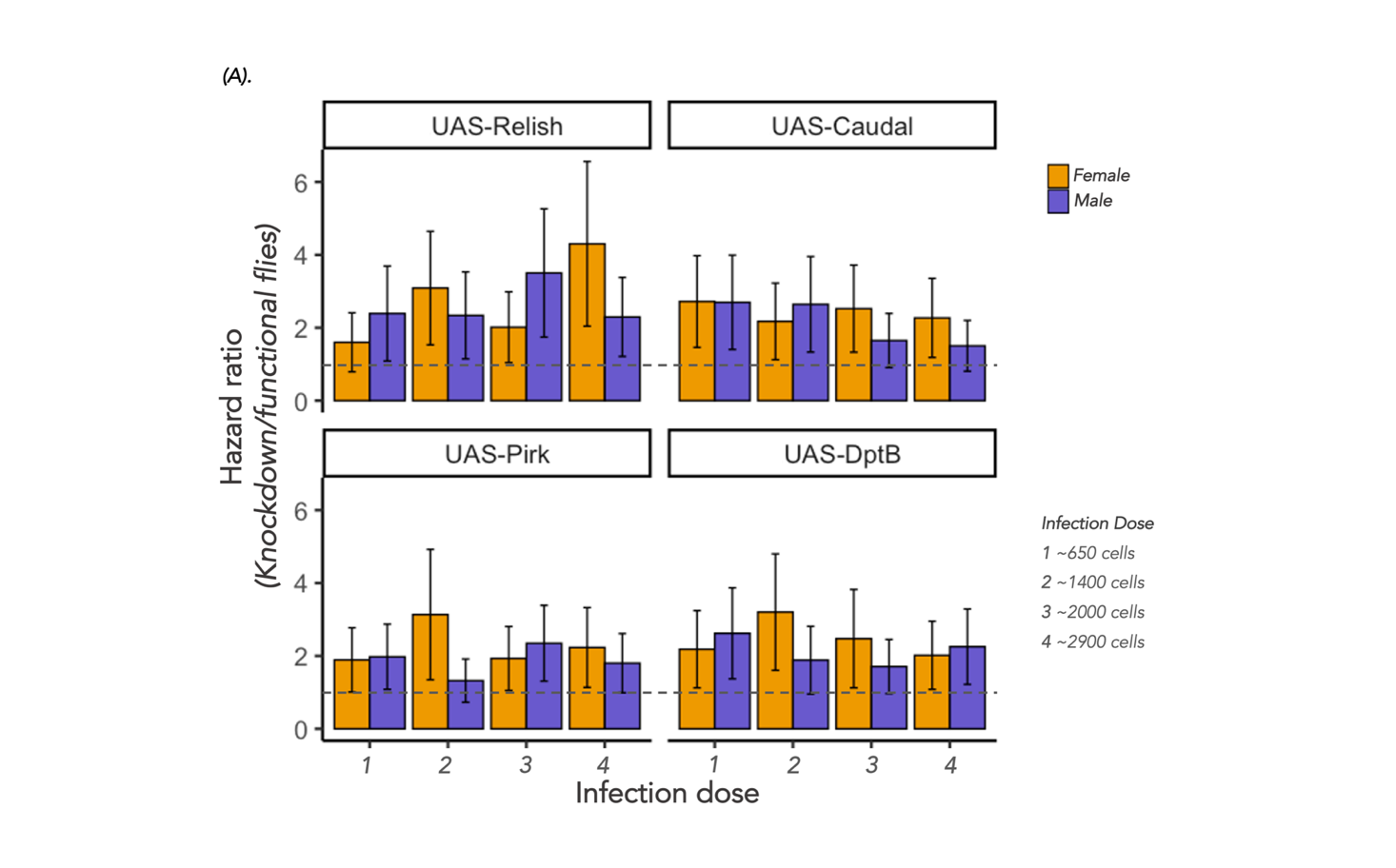

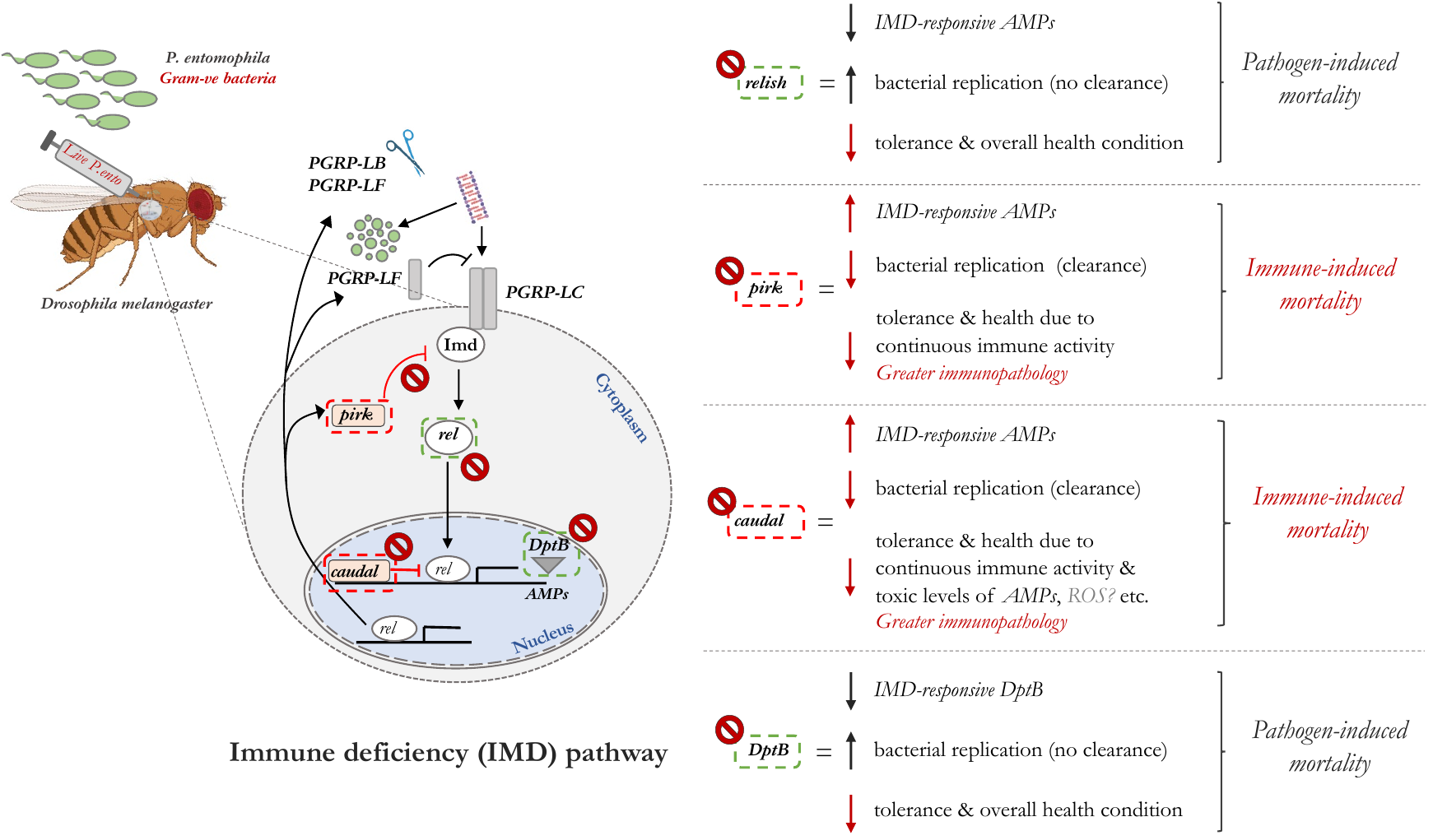
***Figure S6****:* A simplified version of *Drosophila* IMD (immune deficiency) pathway activated by the DAP (diaminopimelic acid) type pedptidoglycan following the gram-negative bacterial infection (*Pseudomonas entomophila*)*.* These receptors then recruit the adaptor molecule IMD and activate the intracellular signalling cascade that leads to the activation of a NF-ĸB *relish*. Upon activation, relish translocates to the nucleus and induces transcription of a set of cocktail of IMD-responsive AMPs such as *Diptericins, Attacins* and *Cecropins*, minutes after pathogen recognition. The negative regulators of IMD such as *pirk, caudal, PGRPs-LB* ensure an appropriate level of immune response, while avoiding immunopathology. Flies with reduced expression of the negative regulators *pirk* and *caudal* show reduced tolerance (overall health state with increasing pathogen burden) despite having higher AMP expression and reducing the bacterial numbers. Flies with reduced *relish* and *DptB* expression also exhibited reduced tolerance and survival mostly due to pathogen-induced mortality because of lower AMPs to target the bacterial replication.

***Table S3:*** **Gene expression of major IMD*-*pathway regulated AMPs**. Gene expression quantified relative to the internal control gene rp49 was quantified in 5-6 replicate individuals (3 pooled flies) of males and females exposed to *P. entomophila* systemic bacterial infection (dose 0.05 OD or ~2000 bac. cells). Analysed using ANOVA and specified the model as gene expression ~ knockdown *_(functional and knockdown)_* x sex *_(male and female)_* x AMPs *_(major IMD-regulated)_* for each of the fly line (functional wild type and IMD-mutants).

| ***Genotype*** | ***Source*** | ***df*** | ***Sum Sq*** | ***F value*** | ***p*** |
| --- | --- | --- | --- | --- | --- |
| *UAS-Relish^RNAi^* | Knockdown *(knockdown and functional flies)* | 1 | 1160.3 | 148.38 | <0.001 |
|  | AMPs *(major IMD-responsive AMPs)* | 2 | 11.2 | 0.717 | 0.49 |
|  | Sex | 1 | 0.4 | 0.051 | 0.82 |
|  | Knockdown x AMPs | 2 | 49.1 | 3.139 | 0.05 |
|  | Knockdown x Sex | 1 | 0.3 | 0.033 | 0.85 |
|  | AMPs x Sex | 2 | 1.4 | 0.087 | 0.91 |
|  | Knockdown x AMPs x Sex | 2 | 11.7 | 0.748 | 0.47 |
| *UAS-Caudal^RNAi^* | Knockdown | 1 | 460.4 | 159.2 | <0.001 |
|  | AMPs | 2 | 10.6 | 1.835 | 0.17 |
|  | Sex | 1 | 23.5 | 8.137 | <0.001 |
|  | Knockdown x AMPs | 2 | 82.3 | 14.24 | <0.001 |
|  | Knockdown x Sex | 1 | 1.1 | 0.394 | 0.53 |
|  | AMPs x Sex | 2 | 4.4 | 0.770 | 0.46 |
|  | Knockdown x AMPs x Sex | 2 | 25.2 | 4.365 | <0.001 |
| *UAS-Pirk^RNAi^* | Knockdown | 1 | 538.3 | 338.9 | <0.001 |
|  | AMPs | 2 | 34.2 | 10.77 | <0.001 |
|  | Sex | 1 | 0.0 | 0.000 | 0.99 |
|  | Knockdown x AMPs | 2 | 47.1 | 14.82 | <0.001 |
|  | Knockdown x Sex | 1 | 8.5 | 5.360 | <0.001 |
|  | AMPs x Sex | 2 | 35.6 | 11.22 | <0.001 |
|  | Knockdown x AMPs x Sex | 2 | 7.2 | 2.255 | 0.11 |

***Table S4:*** **Summary of log_10_ transformed bacterial load** data after exposure to 4 different concentrations (doses) of *P. entomophila* systemic infection, analysed using ANOVA by fitting ‘Knockdown’, ‘Sex’ and ‘dose’ as categorical fixed-effects and replicate ‘vials’ as a random-effects for each of the genotype.

| ***Genotype*** | ***Source*** | ***df*** | ***Sum Sq*** | ***F value*** | ***p*** |
| --- | --- | --- | --- | --- | --- |
| *UAS-Relish^RNAi^* | Knockdown | 3 | 351.07 | 45.4374 | <0.001 |
|  | Sex | 1 | 39.32 | 15.2654 | <0.001 |
|  | Dose | 3 | 518.53 | 67.1097 | <0.001 |
|  | Knockdown x Sex | 1 | 15.53 | 6.0297 | <0.001 |
|  | Knockdown x dose | 3 | 14.43 | 1.8681 | 0.13 |
|  | Sex x dose | 3 | 1.64 | 0.2120 | 0.88 |
|  | Knockdown x Sex x dose | 1 | 3.72 | 1.4450 | 0.23 |
|  | *Random effect* | *Std err* |  |  |  |
|  | *Vials* | *1.605* |  |  |  |
| *UAS-Caudal^RNAi^* | Knockdown | 3 | 157.03 | 20.6574 | <0.001 |
|  | Sex | 1 | 115.72 | 45.6679 | <0.001 |
|  | Dose | 3 | 528.70 | 69.5504 | <0.001 |
|  | Knockdown x Sex | 1 | 0.38 | 0.1497 | 0.69 |
|  | Knockdown x dose | 3 | 27.01 | 3.5529 | 0.014 |
|  | Sex x dose | 3 | 3.55 | 0.4665 | 0.70 |
|  | Knockdown x Sex x dose | 1 | 1.10 | 0.4360 | 0.50 |
|  | *Random effect* | *Std err* |  |  |  |
|  | *vials* | *1.592* |  |  |  |
| *UAS-Pirk^RNAi^* | Knockdown | 3 | 164.02 | 18.3271 | <0.001 |
|  | Sex | 1 | 98.90 | 33.1544 | <0.001 |
|  | Dose | 3 | 407.46 | 45.5296 | <0.001 |
|  | Knockdown x Sex | 1 | 2.57 | 0.8601 | 0.35 |
|  | Knockdown x dose | 3 | 3.17 | 0.3541 | 0.78 |
|  | Sex x dose | 3 | 2.17 | 0.2428 | 0.86 |
|  | Knockdown x Sex x dose | 1 | 0.06 | 0.0215 | 0.88 |
|  | *Random effect* | *Std err* |  |  |  |
|  | *vials* | *1.727* |  |  |  |
| *UAS-DptB^RNAi^* | Knockdown | 3 | 128.94 | 13.2207 | <0.001 |
|  | Sex | 1 | 87.43 | 26.8915 | <0.001 |
|  | Dose | 3 | 615.60 | 63.1179 | <0.001 |
|  | Knockdown x Sex | 1 | 0.44 | 0.1354 | 0.71 |
|  | Knockdown x dose | 3 | 27.20 | 2.7885 | 0.04 |
|  | Sex x dose | 3 | 7.48 | 0.7674 | 0.51 |
|  | Knockdown x Sex x dose | 1 | 1.43 | 0.4411 | 0.50 |
|  | *Random effect* | *Std err* |  |  |  |
|  | *vials* | *1.803* |  |  |  |

***Table S5: Survival analysis.*** Summary of mixed effects Cox model, fitting the model to estimate response to *P. entomophila* infection in flies with fully intact immune system and flies with deficiencies in IMD-pathway. We used data from the individuals of 3-5-day adult males and females infected with 4 concentrations (doses) of *P. entomophila* for each fly lines (IMD-pathway mutants) and specified the model as: survival ~ Knockdown * Sex * dose (1|vial), with ‘Knockdown’ and ‘Sex’ as fixed effects, and ‘vials’ as a random effect. The table shows model output (ANOVA) for survival post-infection for flies with fully function immune system and defects in IMD functioning.

| ***Genotype*** | ***Source*** | ***Loglik*** | ***Chisq*** | ***df*** | ***p*** |
| --- | --- | --- | --- | --- | --- |
| *UAS-Relish^RNAi^* | Knockdown | 1344.5 | 33.429 | 1 | <0.001 |
|  | Sex | 1344.5 | 0.0003 | 1 | 0.98 |
|  | Dose | 1273.7 | 141.55 | 4 | <0.001 |
|  | Knockdown x Sex | 1273.5 | 0.4192 | 1 | 0.51 |
|  | Knockdown x dose | 1270.0 | 7.0930 | 4 | 0.13 |
|  | Sex x dose | 1269.5 | 0.9557 | 4 | 0.91 |
|  | Knockdown x Sex x dose | 1266.8 | 5.4275 | 4 | 0.24 |
|  | *Random effect* | *Std dev* |  |  |  |
|  | *Vials* | *0.244* |  |  |  |
| *UAS-Caudal^RNAi^* | Knockdown | 1337.6 | 28.624 | 1 | <0.001 |
|  | Sex | 1337.5 | 0.0156 | 1 | 0.9 |
|  | Dose | 1285.3 | 104.43 | 4 | <0.001 |
|  | Knockdown x Sex | 1285.3 | 0.0005 | 1 | 0.98 |
|  | Knockdown x dose | 1283.6 | 3.4040 | 4 | 0.49 |
|  | Sex x dose | 1280.6 | 6.0031 | 4 | 0.19 |
|  | Knockdown x Sex x dose | 1279.2 | 2.8850 | 4 | 0.57 |
|  | *Random effect* | *Std dev* |  |  |  |
|  | *vials* | *0.02* |  |  |  |
| *UAS-Pirk^RNAi^* | Knockdown | 1412.1 | 21.126 | 1 | <0.001 |
|  | Sex | 1412.1 | 0.0001 | 1 | 0.99 |
|  | Dose | 1357.4 | 109.30 | 4 | <0.001 |
|  | Knockdown x Sex | 1356.6 | 1.6275 | 1 | 0.20 |
|  | Knockdown x dose | 1355.3 | 2.5330 | 4 | 0.63 |
|  | Sex x dose | 1353.9 | 2.9391 | 4 | 0.56 |
|  | Knockdown x Sex x dose | 1352.3 | 3.2265 | 4 | 0.52 |
|  | *Random effect* | *Std dev* |  |  |  |
|  | *Vials* | *0.14* |  |  |  |
| *UAS-DptB^RNAi^* | Knockdown | 1386.1 | 29.146 | 1 | <0.001 |
|  | Sex | 1386.0 | 0.2848 | 1 | 0.59 |
|  | Dose | 1331.1 | 109.81 | 4 | <0.001 |
|  | Knockdown x Sex | 1331.0 | 0.1187 | 1 | 0.73 |
|  | Knockdown x dose | 1330.8 | 0.3249 | 4 | 0.98 |
|  | Sex x dose | 1329.1 | 3.5419 | 4 | 0.47 |
|  | Knockdown x Sex x dose | 1328.0 | 2.1198 | 4 | 0.71 |
|  | *Random effect* | *Std dev* |  |  |  |
|  | *Vials* | *0.15* |  |  |  |

**Cited references**
